# Biological characteristics of rabbit KLF12 and its regulation of proliferation and apoptosis of ovarian granulosa cells

**DOI:** 10.1101/2025.05.29.656763

**Authors:** Jiawei Cai, Bohao Zhao, Zhiyuan Bao, Yunpeng Li, Xiaoman Han, Yang Chen, Xinsheng Wu

## Abstract

Granulosa cells (GCs), as the largest somatic cell group in follicles, provide nutrients and microenvironment for oocyte growth and maturation. GC proliferation and apoptosis are influenced by several genes, with the Kruppel Like Factor 12 (KLF12) gene contributing to the modulation of several cellular activities. Nevertheless, the precise modulatory function of this gene within the context of ovarian GCs remains unelucidated. We cloned the KLF12 gene of New Zealand female rabbits, and evaluated its expression and localization in the ovaries of female rabbits of different ages by quantitative real-time PCR (qRT-PCR) and fluorescence in situ hybridization (FISH). In order to determine the role of KLF12 in ovarian GCs, we used qRT-PCR, WB, CCK-8 and Annexin V-FITC/PI to detect the effect of KLF12 on GCs proliferation and apoptosis. The coding sequence of KLF12 gene of New Zealand female rabbit was predicted and cloned for the first time in this study. The coding region was 1209 bp, encoding 402 amino acids. KLF12 overexpression has a negative effect on the proliferation and cell cycle progression of rabbit GCs. KLF12 knockdown partially rescues GCs proliferation, accelerates GCs cycle progression, and promotes Estradiol (E_2_) and Progesterone (P) secretion. In addition, KLF12 was also proved to be a key regulator of PI3K/Akt signaling pathway in GCs. In conclusion, the study provides evidence that KLF12 plays a pivotal role in the maturation and proliferation of ovarian GCs in female rabbits. This work establishes a conceptual foundation for investigating deeper into the possible regulatory mechanisms underlying follicular development and growth.

**Author summary:** GCs are essential for oocyte growth, not only providing necessary nutritional support for oocyte growth and maturation, but also constructing a key microenvironment. In this study, the biological function of KLF12 gene was systematically elucidated in rabbit for the first time. It was found that KLF12 inhibited follicular development by negatively regulating GCs proliferation and cell cycle. Combined with its effect on E_2_ and P secretion, the important regulatory role of KLF12 in follicular maturation and hormone balance was revealed. These innovative findings not only deepen the understanding of the regulatory network of ovarian development, but also provide a new theoretical basis for the basic research of reproductive biology. More importantly, it provides a potential molecular target for the development of new technologies to improve the reproductive efficiency of livestock. It also provides a valuable reference model for the study of the mechanism of human reproductive dysfunction related diseases.

## 1. Introduction

As females age, their fertility progressively declines. It is chiefly evidenced by an elevation in the apoptosis of ovarian granulosa cells (GCs), an increase in follicular atrophy, and a substantial reduction in oocyte number and quality, which ultimately depreciates ovarian function(1, 2). Follicles are the primary endocrine and reproductive compartments within the ovary. It essentially consists of oocytes enveloped by adjacent GCs. Being a vital part of the follicular architecture, GCs provide the material support and conducive milieu required for oocyte maturation and dynamic follicular growth. The manifestation of atypical phenomena, including GC apoptosis, can culminate in follicular degeneration and an attenuation of ovarian functionality(3, 4). We previously established an ovarian GC injury model through transcriptome sequencing and found that the Kruppel Like Factor 12 (KLF12) gene is a candidate gene significantly highly expressed during the apoptosis of ovarian GCs(5).

The Kruppel-like transcription factor family is involved in diverse cellular mechanisms, such as cell growth, differentiation, and execution of apoptosis(6). The KLF family members, including KLF4, KLF9, KLF13, and KLF15, play pivotal regulatory role in orchestrating the ovarian function(7, 8). Distinct from other KLF family members, KLF12 is categorized as a zinc finger transcription factor residing within this family. KLF12 restrains the transcription of its target gene through the interaction between its N-terminal PVDLS motif (Pro-Xaa-Asp-Leu-Ser) and the C-terminal binding protein(9, 10). KLF12 is known to activate apoptosis in diverse cell populations, such as endometrial stromal cells(11), ovarian cancer cells(12), and bladder cancer cells(13). Up to now, the action mechanism of the KLF12 gene in ovarian GCs has not been reported.

Our research aims to comprehend the pivotal function of KLF12 in ovarian GC maturation and proliferation. Accordingly, we cloned the full coding sequence (CDS) corresponding to the KLF12 gene from New Zealand female rabbits. Bioinformatic scrutiny of this CDS unveiled insights into the biological role of this gene. Additionally, studies involving overexpression and knockdown have reported that the KLF12 gene may regulate genes related to GC proliferation and apoptosis, including Bax, BCL-2, Caspase-9, and Caspase-3. Simultaneously, KLF12 downregulation activated the PI3K/Akt signaling pathway, modified the expression of key genes involved in E_2_ and P syntheses, and enhanced the release of these hormones by GCs. These findings will aid in obtaining a deeper insight into the role of KLF12 in rabbits.

## 2. Results

### 2.1 Tissue and spatiotemporal expression profiles of KLF12 in female rabbits of different ages

The KLF12 gene expression levels were quantified across various developmental stages in the heart, liver, spleen, lung, kidney, and ovarian tissues obtained from the female rabbits. The KLF12 gene was expressed in different tissues of the rabbits. KLF12 expression was higher in the 12-month-old rabbits than in the 6-month-old rabbits. The KLF12 expression level was the highest in the ovarian tissues, which was significantly higher than that in the 6-month-old rabbits (*P*<0.01) (Figure 1A). Additionally, the results of fluorescence in situ hybridization unveiled that KLF12 gene expression was relatively higher in the GCs of the 12-month-old rabbits (Figure 1B). In summary, KLF12 gene expression in these tissues increased with age, which indicated that this gene plays a crucial biological function in the ovarian tissues of the female rabbits.

**Figure 1.**
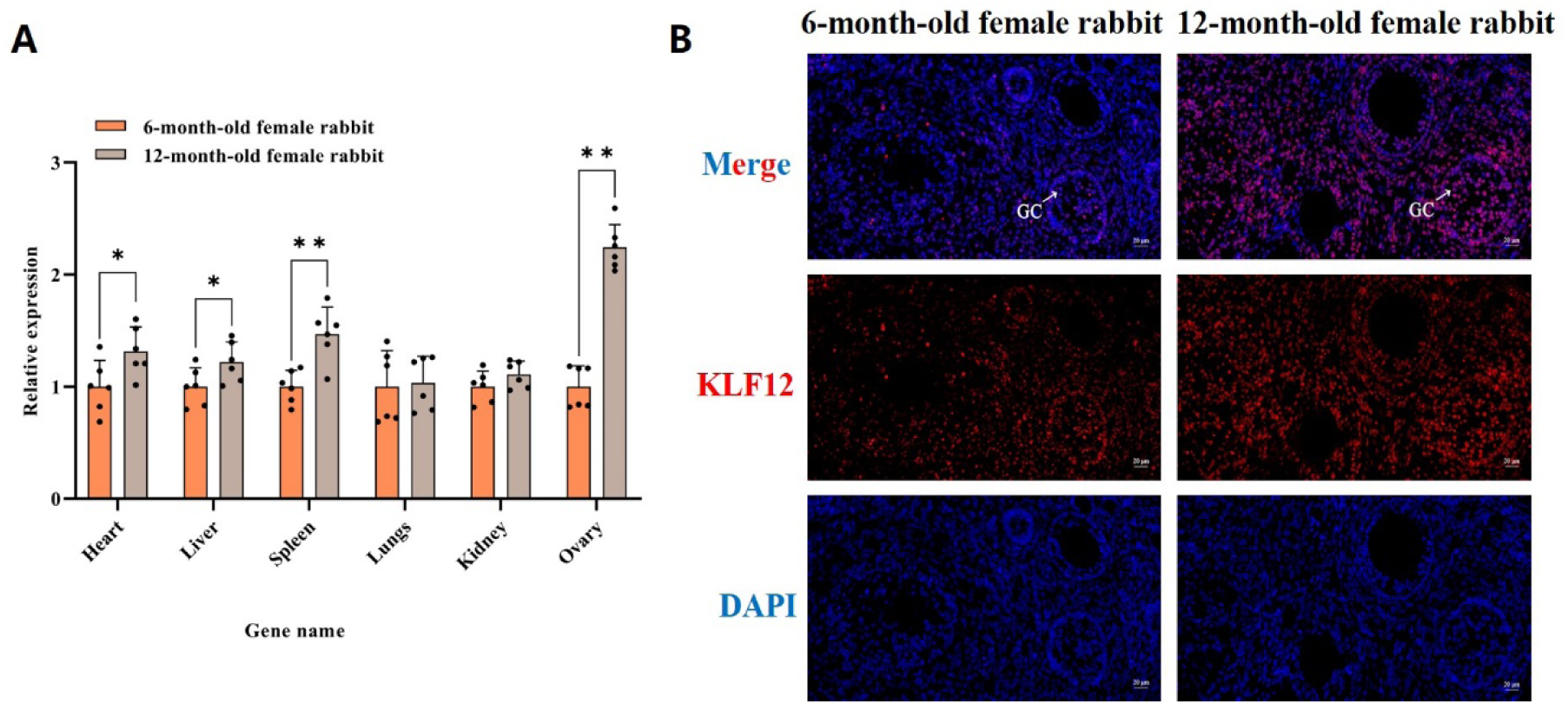
Tissue and spatiotemporal expression profiles of KLF12 in female rabbits of different ages. (A) The KLF12 gene expression level in different tissues of female rabbits at different stages. (B) KLF12 localization in the ovaries of female rabbits. Red denotes KLF12 detected using the fluorescence in situ hybridization probe, and blue indicates the DAPI-stained nucleus. GC, granule cell; Scale bar, 20μm. Data are presented as the mean±SEM (\**P*<0.05, ** *P*<0.01). A two-tailed paired t-test was used for data analyses.

### 2.2 Cloning and bioinformatics analysis of KLF12

Using cloning and sequencing techniques, we here retrieved a 1209-bp-long open reading frame (ORF). This particular ORF encoded a complete protein sequence comprising 402 amino acids, which collectively formed a molecule of approximately 44kDa.

Based on the results of the analysis conducted using the ProtParam tool, the KLF12 protein has a molecular weight of 44,204.83. It has a molecular structure represented by the formula C_1894_H_3065_N_587_O_598_S_18_. The theoretical isoelectric point of KLF12 is 9.74, and the total number of positively and negatively charged residues (Arg+Lys and Asp+Glu) is 52 and 36, respectively. The SignalP-4.1 server predicted that KLF12 has no signal peptide (Figure 2A). The TMHMM server predicted that the KLF12 protein lacks a transmembrane domain (Figure 2B). According to the prediction of the SOPMA software, among the total 402 amino acids in the secondary structure of rabbit KLF12, 65 amino acids (16.17%) formed an α-helix, 73 amino acids (18.16%) formed an extended strand, and 264 amino acids (65.67%) formed a random coil (Figure 2C). The tertiary structure of KLF12 was predicted using the SWISS-MODEL (Figure 2D).

**Figure 2.**
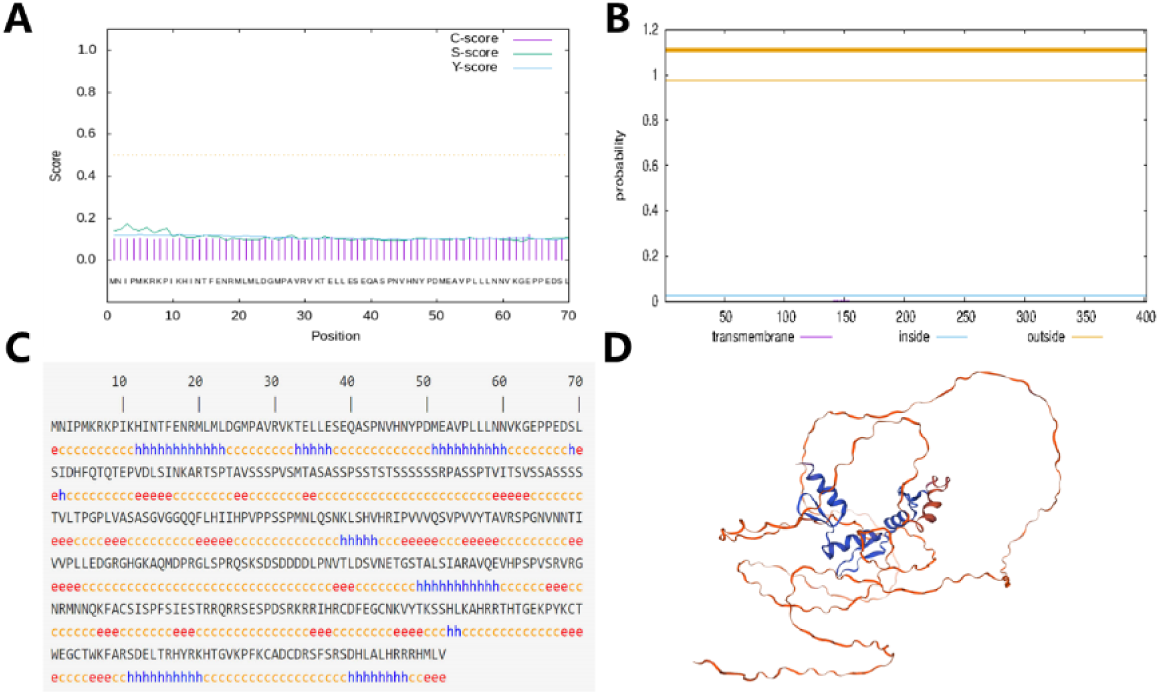
Cloning and bioinformatics analysis of KLF12 (A) SignalP-4.1 software predicted signal peptides of the KLF12 protein. (B) Transmembrane domain prediction of the KLF12 protein. (C) KLF12 secondary structure prediction. h, α-helix; e, extended strand; c, random coil. (D) SWISS-MODEL predicted the KLF12 tertiary structure.

### 2.3 KLF12 plays a negative role in GC proliferation

To determine the role of KLF12 in ovarian GCs, a pcDNA3.1-KLF12 vector was constructed and four siRNAs were designed to knock down KLF12 in rabbit GCs. They were transfected into GCs. The qRT-PCR and WB results unveiled that pcDNA3.1-KLF12 significantly increased KLF12 expression, and siRNA-4 had the most effective knockdown efficiency (*P*<0.01) (Figure 3A).

**Figure 3.**
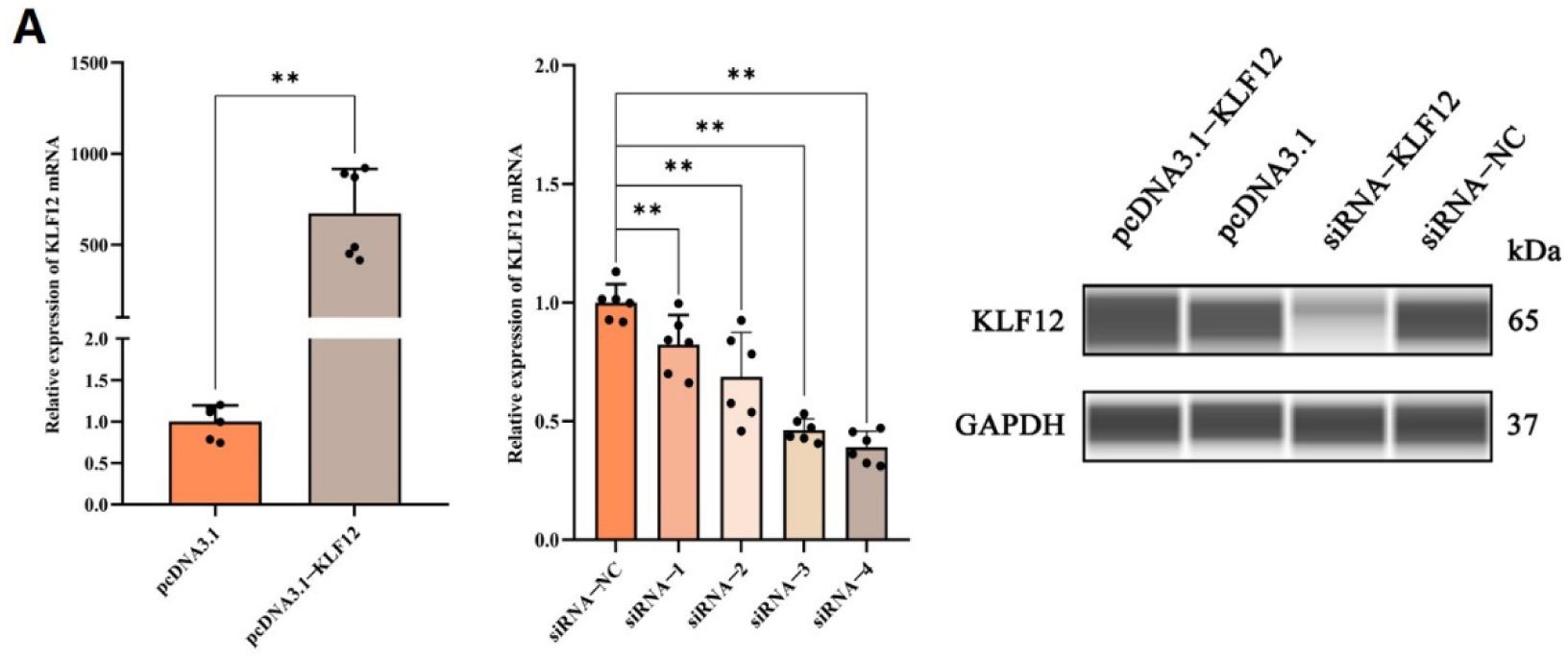
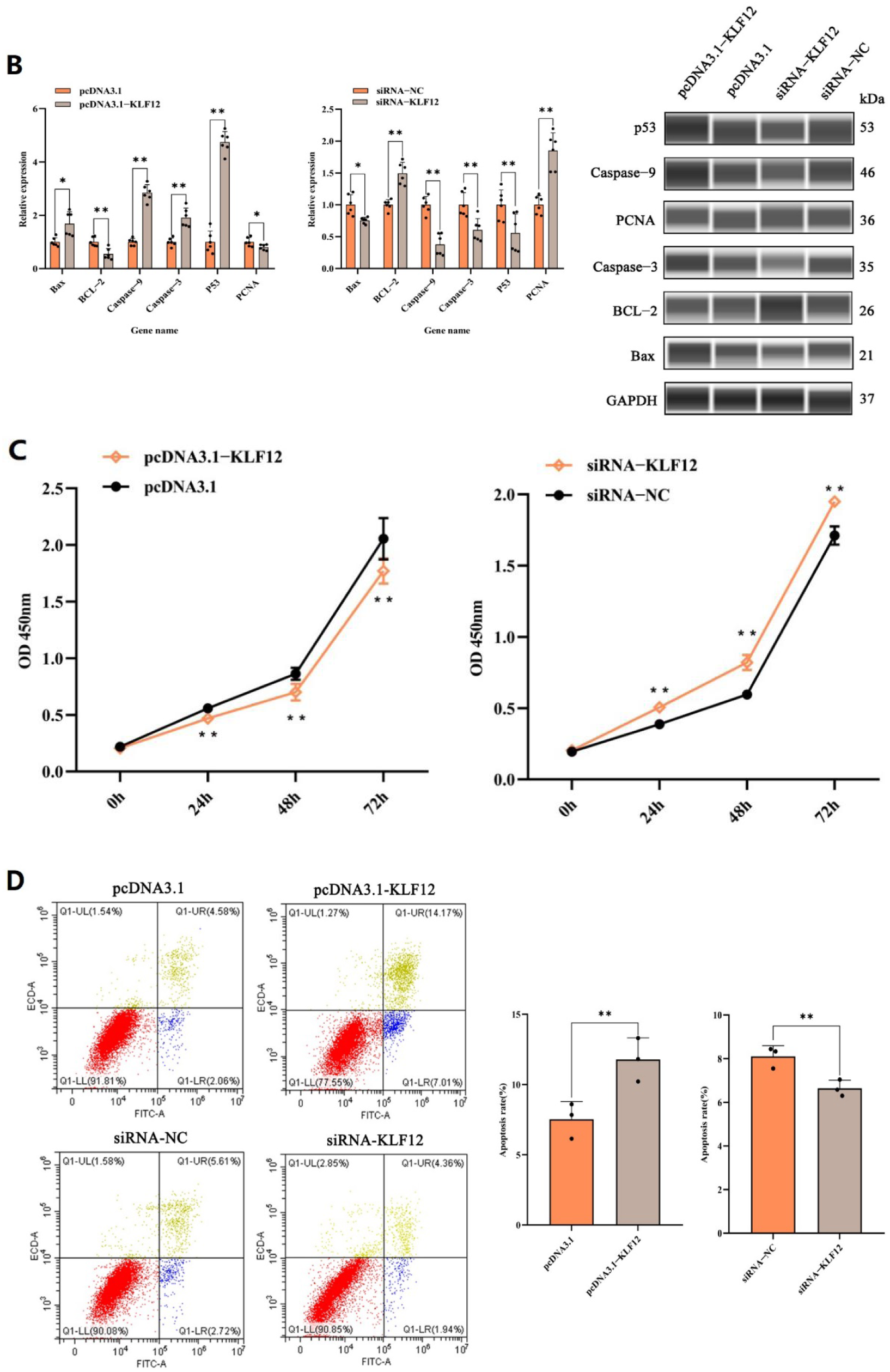
KLF12 plays a negative role in GC proliferation. (A) KLF12 expression was detected after KLF12 overexpression and knockdown in GCs. (B) KLF12 regulated the expression of mRNA and protein related to cell proliferation and apoptosis after KLF12 overexpression and knockdown in GCs. (C) KLF12 inhibited GC proliferation. (D) KLF12 promoted GC apoptosis. Data are presented as the mean±SEM (\**P*<0.05, ** *P*<0.01). A two-tailed paired t-test was used for data analyses.

We detected genes related to GC growth and development. According to the data, KLF12 promoted the mRNA and protein expression of Bax, Caspase-3, Caspase-9, and P53 and inhibited the mRNA and protein expression of BCL-2 and PCNA (*P*<0.05 or *P*<0.01) (Figure 3B). Furthermore, GC proliferation and apoptosis were detected after KLF12 overexpression and knockdown. The findings unveiled that KLF12 notably suppresses GC growth and accelerates their apoptosis (*P*<0.05 or *P*<0.01) (Figure 3C, D).

### 2.4 KLF12 inhibits the expression of cell cycle-related factors in GCs

To study the specific mechanism underlying the promotion of GC proliferation by KLF12 knockdown, qRT-PCR and WB were performed to detect the expression of key cell cycle protein molecules. The mRNA and protein levels of the cell cycle regulatory molecules CCNB1, CCNA2, and CDK4 in KLF12-overexpressing GCs were significantly downregulated compared with those in the empty vector group. By contrast, the CCNB1, CCNA2, and CDK4 expression levels in GCs were significantly increased after KLF12 knockdown (*P*<0.05 or *P*<0.01) (Figure 4A, B).

**Figure 4.**
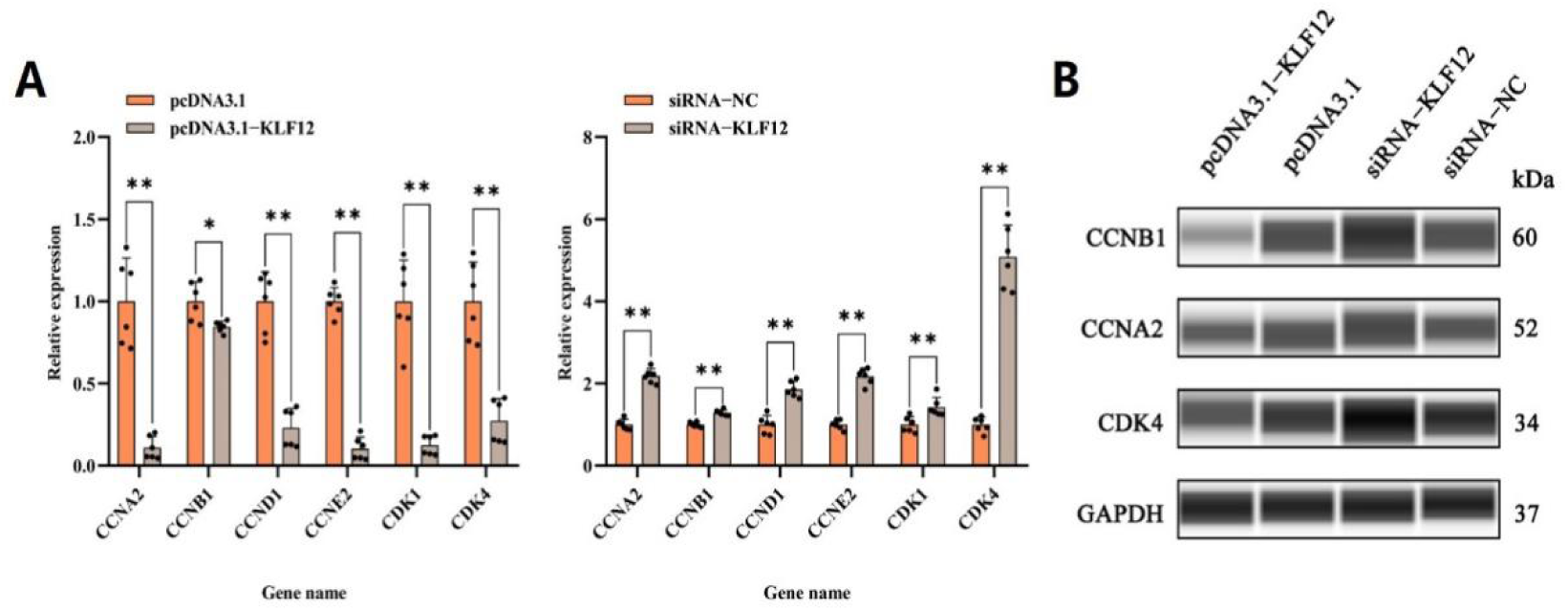
KLF12 inhibits the expression of cell cycle-related factors in GCs. (A) CCNB1, CCNA2, and CDK4 mRNA levels in KLF12-overexpressing GCs and GCs with KLF12 knockdown were detected through qRT-PCR. (B) WB was performed to detect CCNB1, CCNA2, and CDK4 protein levels in KLF12-overexpressing GCs and GCs with KLF12 knockdown. Data are presented as the mean±SEM (\**P*<0.05, ** *P*<0.01). A two-tailed paired t-test was used for data analyses.

The aforementioned studies further demonstrated that KLF12 regulates GC proliferation by regulating the cell cycle.

### 2.5 KLF12 inhibits E_2_ and P levels in GCs

GCs regulate their own proliferation and promote follicular development by secreting hormones, cytokines, etc. Compared with the empty group, E_2_ and P contents secreted by GCs significantly decreased after KLF12 overexpression. By contrast, E_2_ and P contents significantly increased after KLF12 knockdown (*P*<0.01) (Figure 5A, B).

**Figure 5.**
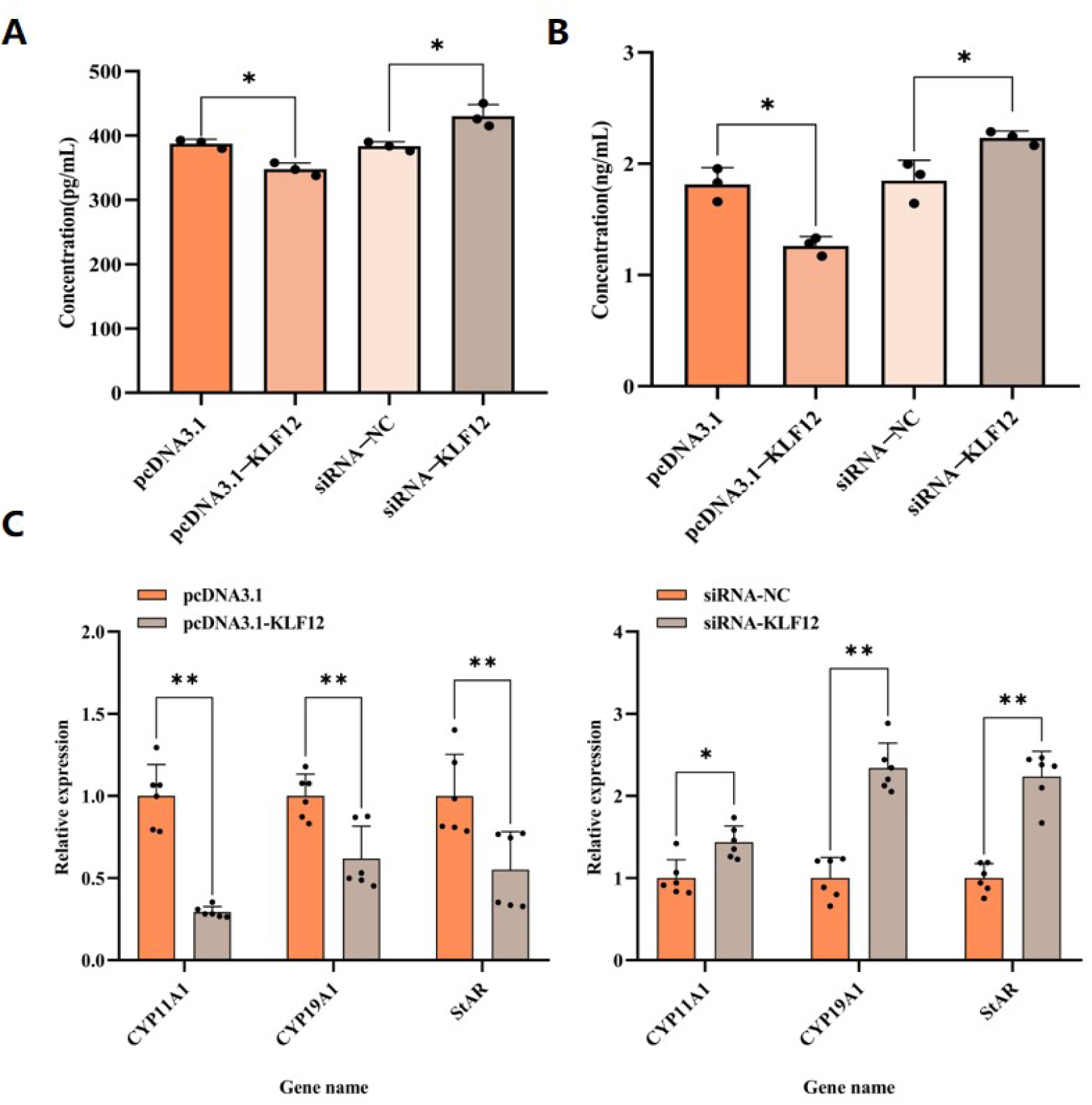
KLF12 inhibits E_2_ and P levels in GCs. (A) E_2_ levels were detected after GCs overexpressed KLF12 or after KLF12 knockdown in GCs. (B) P levels were detected after GCs overexpressed KLF12 or after KLF12 knockdown in GCs. (C) CYP11A1, CYP19A1 and StAR mRNA levels in KLF12-overexpressing GCs and GCs with KLF12 knockdown were detected through qRT-PCR. Data are presented as the mean±SEM (\**P*<0.05, ** *P*<0.01). A two-tailed paired t-test was used for data analyses.

To further explore the reasons for the increased E_2_ and P secretion by GCs after KLF12 knockdown, the mRNA levels of key genes in E_2_ and P syntheses were detected. KLF12 knockdown significantly increased CYP11A1, CYP19A1, and StAR gene expression levels (*P*<0.05 or *P*<0.01) (Figure 5C).

### 2.6 KLF12 regulates the PI3K/Akt signaling pathway in GCs

To delve deeper into the mechanisms underlying KLF12-regulated GC proliferation and apoptosis, we performed WB to quantify the expression levels of key molecules involved in the PI3K/Akt signaling pathway, namely PI3K, phosphorylated PI3K (p-PI3K), Akt, and phosphorylated Akt (p-Akt). The findings revealed that, in contrast to the control (empty) group, the protein levels of the activated forms p-PI3K and p-Akt were notably reduced within the KLF12 overexpression group. Conversely, protein concentrations of these phosphorylated molecules, p-PI3K and p-Akt, were considerably elevated in the KLF12 knockdown group. (Figure 6A, B). The data imply that KLF12 overexpression has a suppressive effect on the PI3K/Akt signaling pathway, whereas KLF12 reduction or silencing leads to a notable stimulation of this pathway.

**Figure 6.**
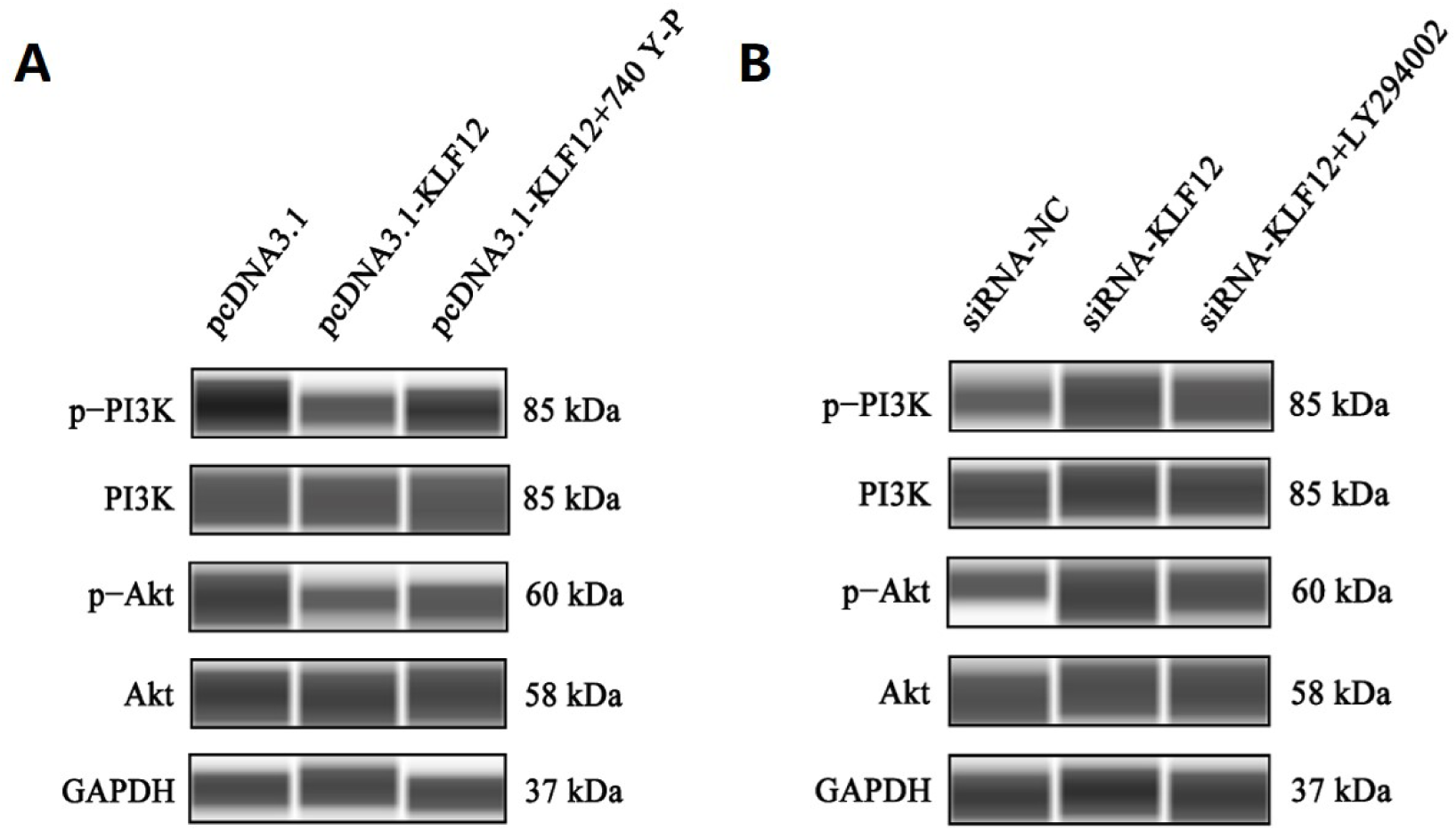
KLF12 regulates the PI3K/Akt signaling pathway in GCs. (A) WB was performed to detect the expression of key molecules in the PI3K/Akt signaling pathway following KLF12 overexpression and 740Y-P addition. (B) WB was performed to detect the expression of key molecules in the PI3K/Akt signaling pathway after KLF12 knockdown and 294002 addition.

To further prove the activation and inhibition effect of KLF12 on the PI3K/Akt signaling pathway of GCs, 740Y-P (10μM) and LY294002 (10μM) were used to treat the transfected GCs. According to the findings, upon the introduction of 740Y-P, p-PI3K and p-Akt protein levels substantially increased within the KLF12-overexpressing group. Conversely, LY294002 addition caused a pronounced decrease in p-PI3K and p-Akt protein concentrations in the cohort with KLF12 knockdown (Figure 6A, B).

The aforementioned outcomes provide additional confirmation that KLF12 indeed acts as a crucial modulator in PI3K/Akt signaling pathway activation within GCs.

## 3. Materials and Methods

### 3.1 Animal and sample collection

Six healthy female New Zealand rabbits aged 6 months (sexually mature, nulliparous) and 12 months (multiparous, parity=4-5) were selected. The selected age groups represented different reproductive stages, and there was no history of infertility or reproductive disorders. Animals were reared under controlled conditions (12:12 light-dark cycle, free use of standard rabbit food and water). The animals were fasted overnight, and their tissue specimens of the heart, liver, spleen, lung, kidney, and ovaries were gathered. The specimens were promptly submerged in liquid nitrogen for instantaneous freezing and transferred to a storage environment maintained at −80℃ for long-term preservation.

Animal experiments were conducted according to the pertinent guidelines for the welfare of laboratory animals, with strict observance of ethical norms and protocols sanctioned by the Animal Care and Use Committee at Yangzhou University (approval No. 202212006).

### 3.2 Cell culture and transfection

Rabbit immortalized GC lines constructed by our group in the early stage were used in this study(14). The GCs were cultured in a DMEM/F12 medium supplemented with 5% fetal bovine serum (FBS) and 1% penicillin/streptomycin solution (Beyotime, Shanghai, China), which is a mixture of 100 units/mL penicillin and 0.1 mg/mL streptomycin.

The cells were cultured in an incubator at 37℃ under 5% CO_2_. The GCs were seeded into a 24-well cell culture dish to ensure 80% cell fusion before transfection was performed using Lipofectamine™ 2000 (Invitrogen, CA, USA). Subsequent experiments were conducted 48 h after transfection.

### 3.3 Bioinformatics analysis of KLF12

The KLF12 coding sequence was analyzed using the DNASTAR software package. ProtParam (http://web.expasy.org/protparam/)(15) was used for predicting the isoelectric point, molecular formula, molecular weight, and instability coefficient of the KLF12-encoded protein. The TMHMM 2.0 (http://www.cbs.dtu.dk/services/TMHMM/)(16) and SignalP 4.1 (http://www.cbs.dtu.dk/services/SignalP-4.1/)(17) online sites were referred to while analyzing proteins for the presence of transmembrane structures and signal peptides. The NPS SOPMA tool was used to predict the protein secondary structure (https://npsa-prabi.ibcp.fr/). The SWISS-MODEL (http://www.swissmodel.expasy.org)(18) was used for predicting the tertiary structure and analyzing the conserved functional domains of their proteins by using the N CBI Conserved Domain Search Service online analysis tool.

### 3.4 KLF12 overexpression and knockdown

To synthesize the overexpression vector for KLF12, high-grade cDNA from the rabbit ovary tissue was synthesized using the PrimeScript™ 1st Strand cDNA Synthesis Kit (Takara, China). The primers were crafted based on the rabbit KLF12 mRNA sequence (GenBank accession number: XM_05185062 1.1) to amplify the KLF12 CDS (primer-F: 5ʹ-gggagacccaagctggctagcATGAATATCCCTATGAAGAGGAAACC-3ʹ, primer-R: 5ʹ-tgctggatatctgcagaattcTCACACCAGCATGTGCCTCC-3ʹ). The amplified CDS was secured through polymerase chain reaction (PCR) by using the Phanta Max Super-Fidelity D NA Polymerase (Vazyme, China) and subsequently inserted into the pcDNA3.1(+) vector. To reduce K LF12 expression, a small interfering RNA (siRNA) was custom designed and synthesized by Shanghai GenePharma. Co., Ltd, and designated as siRNA-1-4. The siRNA sequences are listed in Table 1.

**Table 1.**
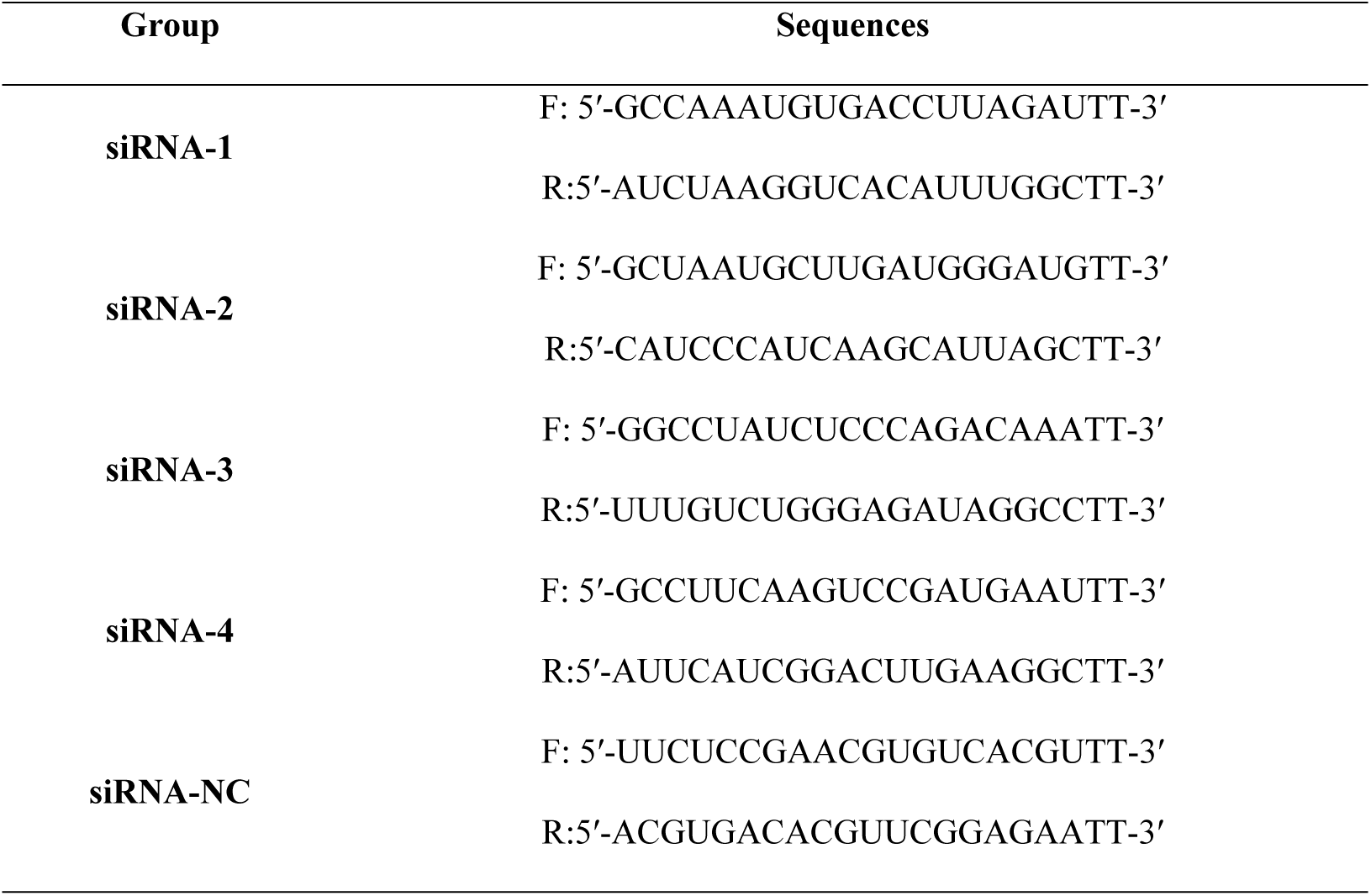
Sequences of siRNA used in the study.

### 3.5 Fluorescence in situ hybridization

The ovarian tissue was fixed with an insitu hybridization fixative for 12h and dehydrated with a series of graded alcohols. Subsequently, the samples were infiltrated with wax, embedded, and sectioned into slices. The sections were immersed in the antigen retrieval solution for 10-15min for heat-induced epitope retrieval. Then, the sections were allowed to cool down to room temperature naturally. Afterward, protease K (20μg/mL) was added to the sections dropwise for digestion at 37°C for 30min. The sections were rinsed with pure water and washed with PBS three times. The pre-hybridization solution was added then added dropwise, and the sections were incubated at 37°C for 1h. The pre-hybridization solution was then poured out, and the 5ng/μL probe-containing hybridization solution (probe synthesis sequence: 5- CY3-TGCGTCCTCCGATGAGCCTTCAGG-3, Wuhan Sevier Biotechnology Co., Ltd.) was added dropwise. Hybridization was performed at 37°C for 12h, and the hybridization solution was washed away. The nucleus was then restained with DAPI, and the antifluorescence quenching sealing agent was added dropwise after washing the sections. The sections were observed under a Nikon positive fluorescence microscope, and the images were captured.

### 3.6 Total RNA extraction, cDNA synthesis, and qRT-PCR

After 48h of transfection, the original medium was discarded, PBS was added to wash the cells, and then 600μL of lysis buffer was added to each well, according to the manufacturer ’s instructions, RNA was extracted from the cells by using the SteadyPure RNA extraction kit. (AG21024; Accurate Biotechnology (Hunan) Co., Ltd., China). Then the concentration of total RNA, 260/230 ratio and 260/280 ratio were evaluated. The ratio was in the range of 2.0-2.1, and the next experiment could be carried out. 500ng RNA was mixed with 2uL gDNA Clean Reaction Mix and 4uL Evo M-MLV RT Reaction Mix, including Evo M-MLV RTase, RNase Inhibitor, dNTPs, Oligo dT (8T) Primer and Random 6 mers Primer, then, RNA was reversely transcribed into cDNA using the Evo M-MLV Mix Kit (AG11728; accurate Biotechnology (Hunan) Co., Ltd.). Gene expression levels were quantified using the SYBR^®^ Green Premix Pro Taq HS qPCR Tracking Kit (AG11733; Accurate Biotechnology (Hunan) Co., Ltd.) on the QuantStudio® 5 system (Applied Biosystems, Thermo Fisher Scientific, Waltham, MA, USA). qRT-PCR was performed using the following conditions: 95°C for 30s, followed by 40 cycles of 95°C for 5 s, 60°C for 30s, and 72°C for 30s. The reaction volume was 20μL, containing 1 μL cDNA, 0.4μL forward and reverse primers (10μM), 10uL 2 ×SYBR^®^ Green Pro Taq HS Premix and 8.2μL RNase free water. Finally, the relative gene expression was measured using the 2^-ΔΔCt^ method(19), and glyceraldehyde 3-phosphate dehydrogenase was used as the internal control. The specific primer sequences are listed in Table 2.

**Table 2.**
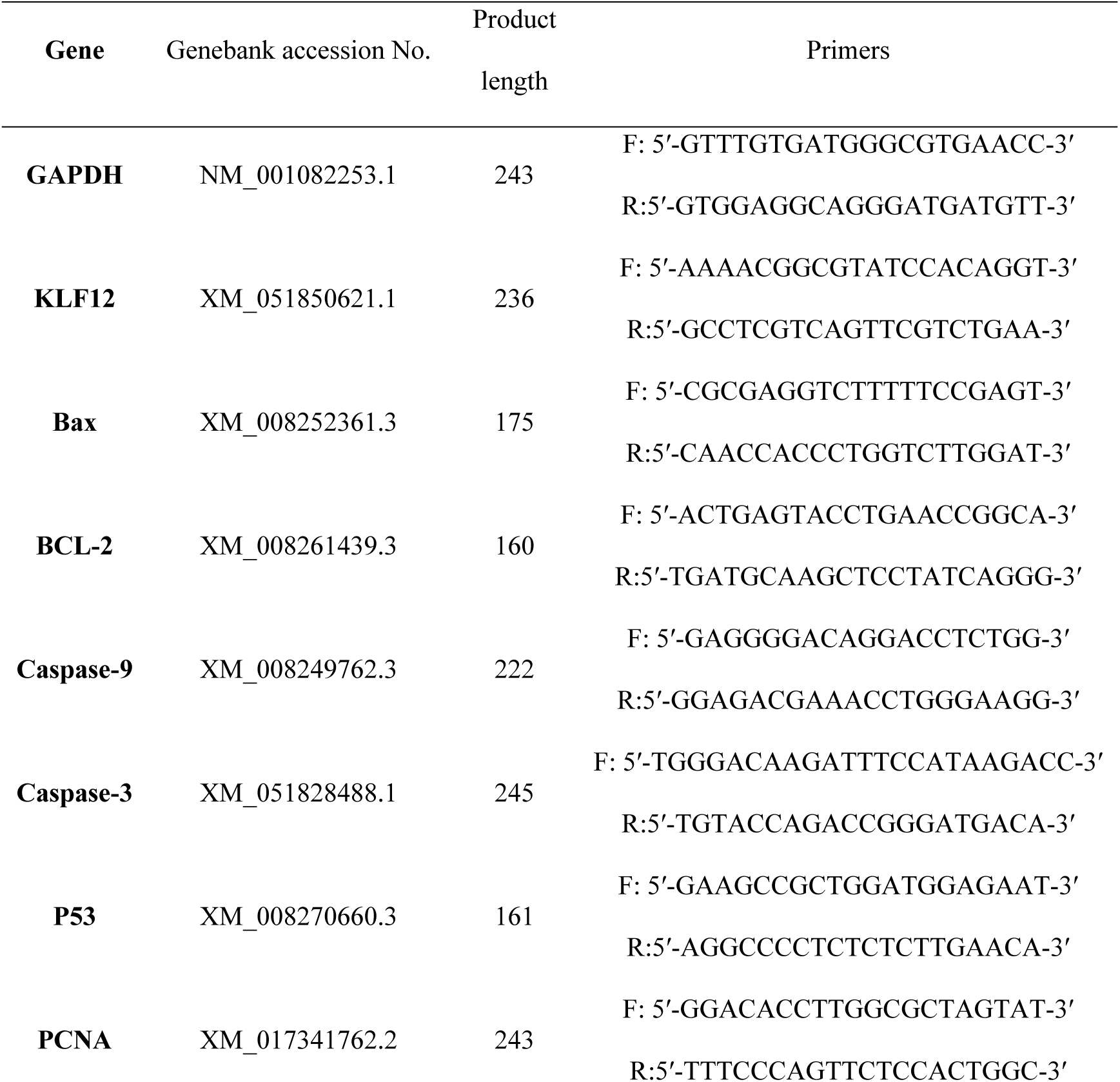
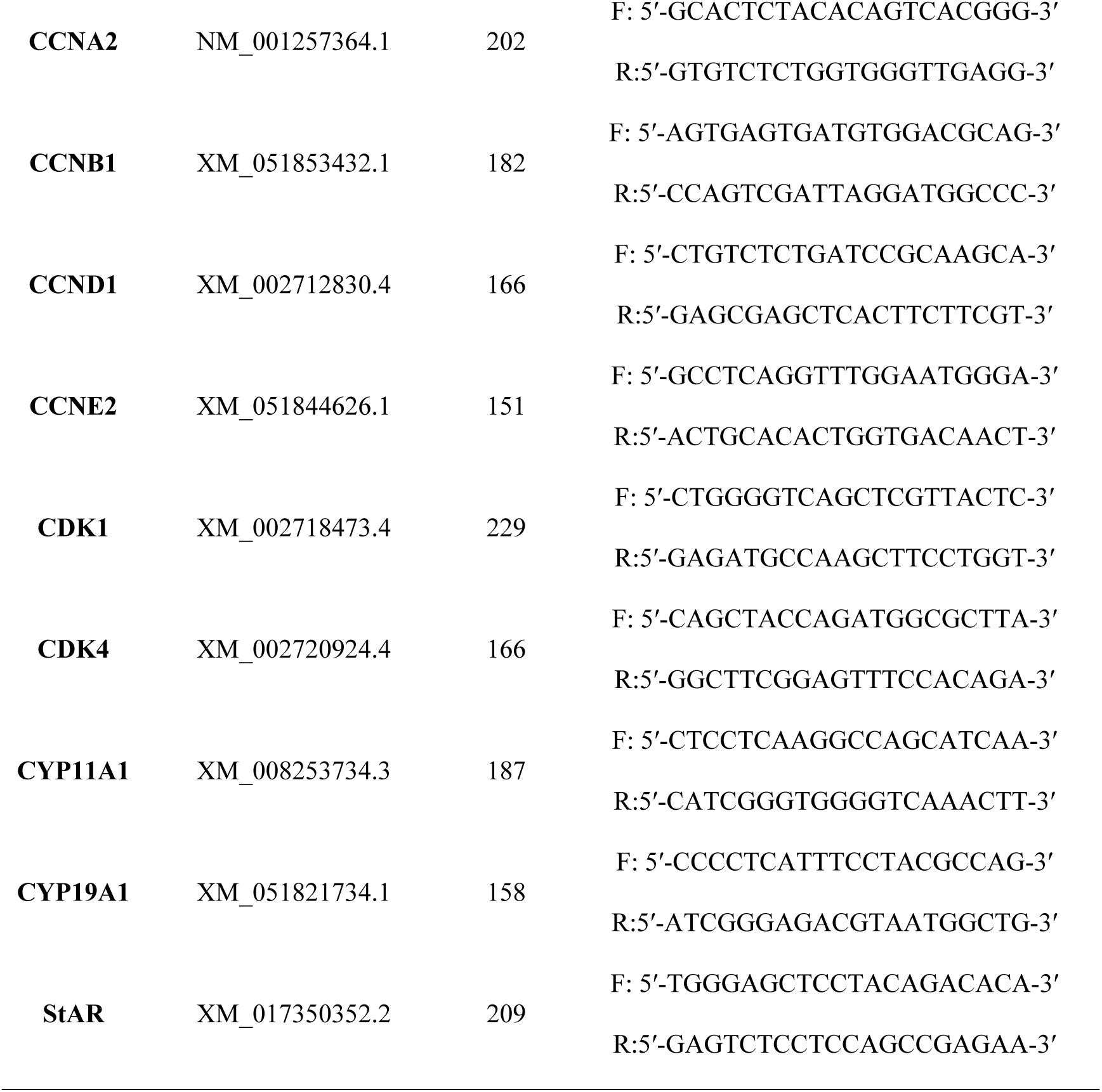
Primer sequences used in qRT-PCR.

### 3.7 Protein extraction and western blotting

The cellular specimens were disrupted using RIPA buffer (Solarbio, Beijing, China) to facilitate protein extraction. The concentration of the extracted proteins was determined using an Enhanced BCA Protein Assay Kit (Beyotime, Shanghai, China). All protein samples were subsequently normalized to a density of 0.5μg/μL, and 3μL of this standardized solution was dispensed into each well. Then, the protein was automatically separated using a western blotting apparatus (ProteinSimple, CA, USA), adhering to the protocol detailed by Harris(20). In all protein analyses, GAPDH was used as an internal reference to ensure standardization. The specific information of the antibodies used in this study is detailed in Table 3, and the original image data of all related experiments are completely saved in the supplementary file.

**Table 3.** The antibodies for Protein Simple Wes Western Blotting System.

| Antibodies names | Source | Dilution Ratio | Vendor | Catalog Number |
| --- | --- | --- | --- | --- |
| **GAPDH** | Mouse | 1:2000 | Proteintech, China | 60004-1-Ig |
| **KLF12** | Rabbit | 1:200 | Proteintech, China | 13156-1-AP |
| **Bax** | Mouse | 1:500 | Proteintech, China | 50599-2-Ig |
| **BCL-2** | Rabbit | 1:500 | Proteintech, China | 26593-1-AP |
| **Caspase-9** | Rabbit | 1:500 | Proteintech, China | 10380-1-AP |
| **Caspase-3** | Rabbit | 1:200 | Proteintech, China | 25128-1-AP |
| **p53** | Mouse | 1:500 | Proteintech, China | 60283-2-Ig |
| **PCNA** | Mouse | 1:500 | Proteintech, China | 60097-1-Ig |
| **CCNB1** | Rabbit | 1:200 | Proteintech, China | 55004-1-AP |
| **CCNA2** | Mouse | 1:500 | Proteintech, China | 18202-1-AP |
| **CDK4** | Rabbit | 1:500 | Proteintech, China | 11026-1-AP |
| **PI3K** | Rabbit | 1:200 | Proteintech, China | 60225-1-Ig |
| **Akt** | Mouse | 1:200 | Proteintech, China | 60203-2-Ig |
| **p-PI3K** | Mouse | 1:200 | Affinity, China | AF3241 |
| **p-Akt** | Mouse | 1:200 | Proteintech, China | 66444-1-Ig |

### 3.8 Cell apoptosis and proliferation assay

Cellular apoptosis was analyzed using the Annexin V-FITC Apoptosis Detection Kit (Vazyme, China). Apoptosis levels were quantified using a flow cytometer (FACSAria SORP model; Becton Dickinson, USA). Following acquisition, the cytometric data were analyzed using FlowJo V10 software (FlowJo, OH, USA). Meanwhile, cell proliferation was detected using the cell Counting Kit-8 (Vazyme, China). The optical density values of 96 holes at 0, 24, 48, and 72h were measured at 450nm by using Infinite M200 pro (Tecan, Switzerland).

### 3.9 Enzyme-linked immunosorbent assay

Enzyme-linked immunosorbent assay (ELISA) kits for E_2_ (Cat.# ml027864, the minimum detectable concentration was less than 1.0pg/mL, linear range: 15-480pg/mL) and P (Cat.# ml041624, the minimum detectable concentration was less than 0.1ng/mL, linear range: 0.25-8g/mL) were obtained from Mlbio (Shanghai, China). According to the manufacturer’s instructions, E_2_ and P levels in the cell culture supernatant were measured using the ELISA kits.

### 3.10 Statistical analysis

Statistical significance was determined by two-tailed Student ’s t test or one-way analysis of variance (ANOVA) in SPSS 25.0 statistical software package (SPSS Inc., Chicago, IL, USA). All collected data were then expressed as mean±standard error (SEM). In order to meet the normality hypothesis, the Shapiro-Wilk test was used to verify the normal distribution of the data, and Levene ’s Test was used to verify the homogeneity of the variance. After performing analysis of variance, we use Tukey ’s HSD multiple comparison test method to ensure the significance of the differences between groups. Furthermore, the data were graphically represented using GraphPad Prism 8 (GraphPad Software, Inc., San Diego, CA, USA). Results were considered statistically significant at *P*<0.05 (*) and highly significant at *P*<0.01 (**).

## 4. Discussion

GCs are the largest somatic cell population in the ovary. They provide material guarantee and a microenvironment for oocyte maturation and follicular development. During follicular growth and development, only a very small number of follicles successfully develop and ovulate, whereas most of the remaining follicles undergo atresia. The key cellular event mediating this physiological process is GC apoptosis. We previously found that KLF12, a member of the Kruppel-like factors family, was upregulated during ovarian GC apoptosis. In the present study(5), the KLF12 expression level was higher in 12-month-old female rabbits than in 6-month-old female rabbits, and the expression level was the highest in the ovarian tissue. Subsequently, we cloned the coding sequence of the rabbit KLF12 gene, which encodes a protein composed of 402 amino acids. A bioinformatics evaluation revealed that the KLF12 protein has favorable stability characteristics, lacks a signal peptide, and does not feature a transmembrane domain.

To regulate gene transcription, KLFs bind to the GC box or CACCC of the target gene promoter region through a highly conserved carboxyl-terminal DNA domain, thereby affecting cell proliferation, differentiation, and apoptosis(9, 21). KLF9 levels are meticulously controlled to preserve the cellular equilibrium, which encompasses processes such as cell proliferation, extracellular matrix formation, cell migration, and adherence(22, 23, 24, 25, 26). KLF5 is involved in promoting the ovarian cancer stem cell-like phenotype(27). KLF8 promotes the induction of tumorigenic breast stem cells by targeting miR-146a(28). KLF4, KLF9, and KLF13 transcripts are expressed in ovarian cells and regulate these cells(8). These collective results indicate that KLFs are promising therapeutic targets in modulating cell proliferation and apoptosis. Intriguingly, our study revealed that heightened KLF12 expression concurrently facilitates apoptosis induction in ovarian GCs. KLF12 gene knockdown and overexpression can regulate the expression of GC proliferation- and apoptosis-related genes Bax, BCL-2, Caspase-9, and Caspase-3, and can affect the expression of cell cycle-related proteins CCNB1, CCNA2, and CDK4. Cell cycle regulation is a highly fine-tuned process, which results in cell growth, genetic material replication, and cell division. The PI3K/Akt signaling pathway can activate Cyclin A, Cyclin B, Cyclin D, and Cyclin E, to promote GC proliferation(29).

The PI3K/Akt pathway, serving as a significant axis governing cell proliferation and apoptosis, plays a critical role in orchestrating GC growth and programmed cell death throughout follicular maturation(30, 31, 32). PI3K is activated when growth factors and cytokines bind to their cell surface receptors(33). As the main target of proto-oncogenes and PI3K, Akt regulates cell survival and proliferation(34). It oversees cellular proliferation and protein synthesis and counteracts pro-apoptotic entities such as BAD and Caspase-9, which are pivotal elements in executing programmed cell death. Within the scope of this investigation, we observed that KLF12 overexpression notably reduced p-PI3K and p-Akt protein levels. Conversely, when KLF12 was knocked down, p-PI3K and p-Akt protein concentrations were significantly elevated. Subsequently, the transfected GCs were treated with 740Y-P and LY294002. After 740Y-P addition, p-PI3K and p-Akt protein levels significantly increased in the KLF12 overexpression group, whereas the protein levels of the phosphorylated molecules p-PI3K and p-Akt significantly decreased after LY294002 addition. This result again proves that KLF12 molecules exert a regulatory effect on the PI3K/Akt signaling pathway.

In addition, as the main unit of follicles, GCs are the key to ensure animal oocyte maturation and maintain normal hormone levels. The initial stage in P synthesis is the translocation of cholesterol from the exterior to the interior of the mitochondrial membrane. StAR, a pivotal component, governs steroid production. Then, CYP11A1 transforms cholesterol into pregnenolone, and mitochondrial enzymes-mediate the subsequent conversion of pregnenolone into progesterone(35). KLF12 knockdown changes the expression of key genes involved in E_2_ and P syntheses and promotes E_2_ and P secretion by GCs. This indicates that KLF12 can regulate E_2_ and P levels by regulating the key proteins involved in their syntheses.

## 5. Conclusion

In conclusion, the coding sequence of KLF12 gene of New Zealand female rabbit was predicted and cloned for the first time in this study. The coding region was 1209 bp, encoding 402 amino acids. KLF12 overexpression has a negative effect on the proliferation and cell cycle progression of rabbit GCs. KLF12 knockdown partially rescues GCs proliferation, accelerates GCs cycle progression, and promotes E_2_ and P secretion. In addition, KLF12 was also proved to be a key regulator of PI3K/Akt signaling pathway in GCs. The results provide a theoretical basis and experimental evidence for the future study of the regulatory role of KLF12 in ovarian development of New Zealand female rabbits, and lay a conceptual foundation for further study of the possible regulatory mechanisms of follicular development and growth (Figure 7).

**Figure 7.**
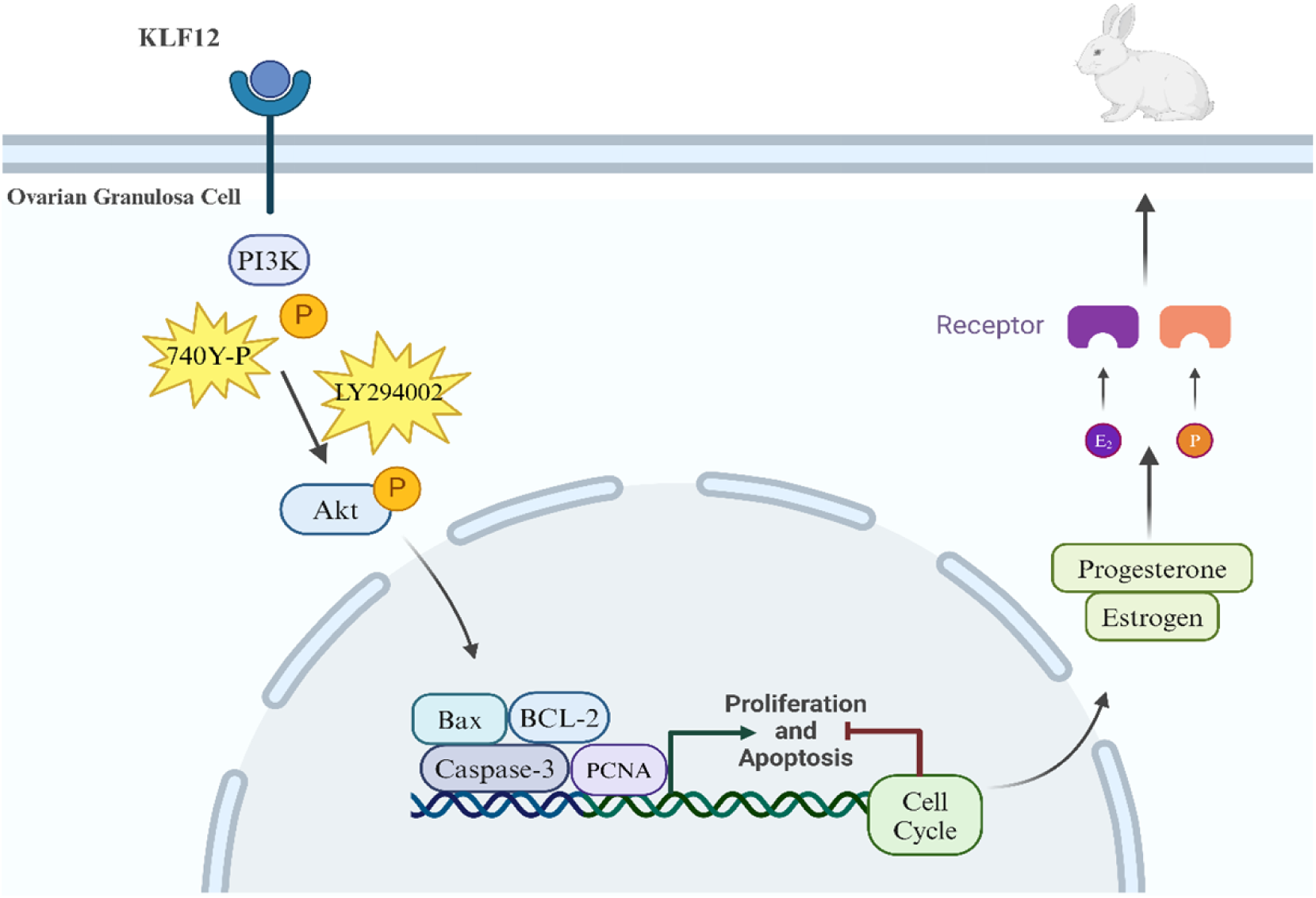
The mechanism of KLF12 regulating the proliferation and apoptosis of rabbit ovarian granulosa cells.

## Conflicts of Interest

We certify that there is no conflict of interest with any financial organization regarding the material discussed in the manuscript.

### Author Contributions

Jiawei Cai: Writing-review & editing, Writing-original draft. Bohao Zhao: Writing-review & editing. Zhiyuan Bao: Resources, Methodology. Yunpeng Li: Resources, Methodology. Xiaoman Han: Formal analysis. Yang Chen: Writing-review & editing. Xinsheng Wu: Visualization, Supervision, Project administration.

## Acknowledgments

This research was funded by the China Agriculture Research System of MOF and MARA (CARS-43-A-1).

